# An Atomic Force Microscopic analysis of Exosomes derived from Tumor Associated pluripotent Mesenchymal Stem Cells

**DOI:** 10.1101/2025.07.18.665627

**Authors:** Tanmay Kulkarni, Simone Newell, Erik Armstrong, Narendra Banerjee, Jasmine Cuffee, Santanu Bhattacharya, Hirendra Banerjee

**Author notes:** Corresponding author address: Santanu Bhattacharya, Griffin 412, Mayo Clinic, 4500 San Pablo Rd, Jacksonville, FL, 32224, USA, Hirendra Banerjee, Room#425, Elizabeth City State University, 1704 Weeksville Rd, Elizabeth City, NC, 27909, USA, Corresponding author email address.

## Abstract

Mesenchymal stem cell (MSC)-derived exosomes are recognized as critical mediators within the tumor microenvironment (TME), exerting both pro- and anti-tumorigenic effects depending on contextual factors. These vesicles are also gaining attention for their potential as therapeutic vehicles in regenerative medicine and targeted drug delivery. However, the influence of the TME on the physical characteristics of MSC-derived exosomes remains poorly understood. In this study, we utilized Atomic Force Microscopy (AFM) to investigate the morphological and nanomechanical properties of MSC-derived exosomes under standard and TME-like conditions. AFM imaging in fluid mode preserved the native structure of exosomes and enabled high-resolution analysis of their topography, surface roughness, stiffness, adhesion, and deformation. AFM offers unique advantages in exosome research by enabling label-free, nanoscale analysis of vesicle properties in near-physiological conditions. The ability to detect such subtle but functionally significant changes highlights the relevance of AFM in exosome characterization and quality assessment. Our results revealed that exposure to the TME induces marked changes in exosomal membrane morphology and mechanical behavior, including increased surface heterogeneity, higher stiffness, and altered adhesive interactions. These biophysical alterations may reflect changes in membrane composition and protein or lipid cargo, potentially affecting exosome function, uptake, and therapeutic efficacy. Overall, our findings provide new insights into how the TME modulates MSC exosome biophysics and underscore the utility of AFM-based techniques for advancing the development of exosome-based diagnostics and therapies.

## Introduction

The tumor microenvironment (TME) is a complex and dynamic niche composed of a heterogeneous population of cells, signaling molecules, vasculature, and the extracellular matrix (ECM) [1, 2]. Among the non-malignant cells present in the TME, mesenchymal stem cells (MSCs) play a pivotal role in modulating tumor progression[3]. A key mechanism through which MSCs influence the TME is via the secretion of exosomes, nano-scale vesicles loaded with bio-molecular cargo such as RNA, DNA, and proteins[4, 5]. These exosomes function as critical messengers, orchestrating a variety of cellular processes that promote tumor growth and metastasis[6, 7].

MSCs are multipotent stromal cells that can differentiate into various cell types, including osteoblasts, chondrocytes, and adipocytes[8]. In the context of cancer, they are integral components of the TME, contributing to the formation of tumor stroma and facilitating the anchorage of tumor cells[9]. Moreover, MSCs can transdifferentiate into other cell types such as M2-type macrophages and myeloid-derived suppressor cells (MDSCs), which further enhance the pro-tumorigenic milieu[10, 11].

Exosomes have emerged as a promising platform for drug delivery due to their unique properties and functional characteristics[12, 13]. These nanosized extracellular vesicles, typically ranging from 30 to 150 nm in diameter, demonstrate significant advantages over traditional drug delivery systems such as liposomes and synthetic nanoparticles [14]. Their innate features include low toxicity, high biocompatibility, and the ability to cross biological barriers, including the crucial blood-brain barrier (BBB)[15, 16].

Among the various sources of exosomes, those secreted by mesenchymal stem cells are gaining increasing attention for their roles in modulating the TME [5, 17]. These vesicles are capable of delivering diverse molecular cargo directly into recipient cells, thereby influencing key cellular pathways and promoting tumor progression. Emerging evidence suggests that these exosomes play a significant role in the suppression of anti-tumor immunity and the development of resistance to cancer therapies [18, 19]. This indicates their pathological involvement in tumor growth and progression.

Atomic Force Microscopy (AFM) is a versatile and powerful tool that has found widespread application in biological research due to its capacity for high-resolution imaging, nanomechanical characterization, and operation in near-physiological environments [20]. Unlike electron microscopy, which often requires extensive sample preparation, AFM enables label-free, real-time imaging of biological samples under native conditions, preserving structural integrity and functionality [21]. In cellular and molecular biology, AFM has been extensively employed to probe the surface topography and mechanical properties of various biomaterials, including cells, membranes, proteins, and exosomes [22-28]. Its ability to perform force spectroscopy allows precise quantification of nanomechanical parameters such as stiffness (Young’s modulus), adhesion, and deformation, which are critical indicators of physiological or pathological changes at the nanoscale [29]. AFM enables the visualization of exosomal morphology with nanometer resolution, distinguishing subtle alterations that might be undetectable by techniques like nanoparticle tracking analysis (NTA) or dynamic light scattering (DLS) and transmission electron microscopy (TEM). Furthermore, AFM-based nanomechanical profiling can reveal changes in exosome stiffness and elasticity, which are associated with disease progression, including cancer [30, 31].

Recent studies have demonstrated that tumor-derived or tumor-exposed exosomes exhibit distinct mechanical properties compared to those from healthy or unexposed cells. These nanomechanical signatures, measurable by AFM, may serve as potential biomarkers for diagnosis or therapeutic targeting [32]. The combination of AFM imaging and force spectroscopy thus provides a comprehensive biophysical characterization of exosomes, facilitating a deeper understanding of their roles in the tumor microenvironment (TME) and their impact on cellular behavior.

In this study, we utilized AFM to investigate the morphological and nanomechanical differences between exosomes derived from Mesenchymal Stem Cells (MSCs) exposed to the tumor microenvironment and those from unexposed MSCs. This approach aims to elucidate how the TME influences exosomal properties and contributes to tumor-associated signaling.

## Materials and Methods

### Cell Culture

NCl-H209 (ATCC HTB-172), Bone marrow-derived mesenchymal stem cells (ATCC PCS-500-12) were acquired from American Type Culture Collection (ATCC, Manassas, Virginia). Routine maintenance for each cell line was followed as per ATCC protocol. All media were supplemented with 10% fetal bovine serum and 100 µg/mL penicillin/streptomycin. All cell lines were grown in 5% CO^2^ in 25 cm^2^ filter cap flasks and cultured in a laboratory incubator at 37°C.

### MSC-TME Exposure

In six-well plates, MSC cells were grown to confluence, placed under inserts with 0.4 µm micropores, which contained HTB172 lung cancer cells grown to confluence along with controls with no cancer cells. These cells were cultured for a period of 48 hours before exosomes were isolated.

### Exosome Isolation

Exosomes were isolated following the total exosome isolation kit purchased from Invitrogen, Carlsbad, CA (Catalog # 4478359).As per the manufacturer’s protocol, from a confluent flask of cells, culture media was collected in a 15 ml centrifuge tube and centrifuged at 14,000 rpm, 10 min. Supernatant was discarded and the pellet was re-suspended in 5 mL of appropriate incomplete media (no Fetal Bovine Serum), placed in a 25 cm^2^ culture flask, and allowed to grow for up to 12 hrs. This was done to eliminate the high levels of exosomes contained in FBS and to prevent contamination of the cell derived exosomes. After growing cells in incomplete media for 10 h, cells in media were collected into a 15 ml centrifuge tube and centrifuged at 14,000 rpm for 10 min. Supernatant (1 mL) was placed in each of 5 Eppendorf tubes (1.5 mL). Supernatant was then centrifuged at 2,000 x g for 30 min to remove cells and debris and supernatant was then transferred from each tube and placed into a new 1.5 mL Eppendorf tube without disturbing the pellet (total 5 tubes). Total Exosome Isolation reagent (0.5 volumes) was added to each of the 5 tubes containing cell-free culture media. The culture media/reagent mixture was vortexed until a homogenous solution appeared and the mixture was incubated overnight at 4°C. After incubation, the samples were centrifuged at 10,000 x g for 1 hour at 4°C in the benchtop Centrifuge 5804 R (Eppendorf, Hauppauge, NY). The supernatant was aspirated and discarded leaving the pellet containing exosomes, which was not readily visible in most cases. The pellet was then resuspended in 1 mL 1X PBS and stored in minus eighty degree Celsius freezer.

### AFM experimental methodology

Freshly cleaved mica surface, which is considered to be optimally flat and possesses a slight negative charge, was employed to study the morphology and nanomechanical attributes of exosomes under various treatments. To achieve an effective adsorption of exosomes on mica surface, freshly cleaved mica was modified using a 3:1 mixture by volume comprising of 3-aminopropyltriethoxysilane (APTES) and N, N-diisopropylethylamine (DIPEA) for 2 hours at 60 °C [15]. The modified mica surface was used immediately. For AFM experiments, the exosome samples were diluted 1000-fold with Milli Q water. Working exosome sample was prepared by diluting the original exosome stock solution 1000-fold with Milli Q water. Following which, 5 µL of working solution was drop-casted on the modified mica to let the exosomes adhere for 30 mins. Mica surface was then rinsed with Milli Q water to remove any unbound exosomes. To conduct AFM experiments, we employed Dimension Icon Scanasyst AFM (Bruker Corporation, Santa Barbara, CA). To evaluate the surface morphology and nanomechanical attributes of the exosomes, we incorporated Peak Force Quantitative Nanomechanical Mapping (PF-QNM) and Nanoindentation experimental techniques, respectively. PFQNM, a specialized tapping mode generates a height image and quantitates nanomechanical attribute of the sample simultaneously. In Peak force tapping mode, where the AFM tip intermittently meets the sample surface to generate a feedback signal yielding a surface topography. A Scanasyst Air probe bearing a sharp tip has a radius of 5 nm and a stiff cantilever with nominal spring constant of 0.4 N/m and with pyramidal tip geometry was used to achieve high resolution topography of the exosomes. For characterization of topographical morphology, the laser was aligned on the back of the cantilever (gold reflective surface) to achieve an optimal signal-to-noise ratio. Topography was assessed using a low peak force of 300 pN and scan rate of 0.1 Hz. At least 50 samples were analyzed to evaluate the surface topography of the exosomes. Nanoscope Analysis v1.9 software was then employed to analyze morphological characteristics such as the height and surface roughness of the exosomes. The height of the exosomes was analyzed using the sectional analysis module under the analysis software whereas, the surface roughness was derived as the root mean square variation in the feature height, also available as an inbuilt module. For nanomechanical properties characterization, AFM probe was calibrated in the presence of Milli Q water on a freshly cleaved mica surface to determine the actual spring constant of 0.38 N/m and deflection sensitivity of 35 nm/V. The calibration of AFM tip in fluid medium allows compensation for the hydrodynamic drag that exists during the AFM experiments. Nanoindentation experiment was performed on at least 50 samples in which, each tip-sample interaction resulted into a force-separation (F-S) curve. A trigger force of 700 pN was applied to engage the tip with the sample. Further, by employing DMT model, each F-S curve was analyzed to yield various nanomechanical attributes such as Young’s modulus (YM), deformation and adhesion.

## Results

### Morphology of MSC-Derived Exosomes Under Various Conditions

Mica, being optimally flat with a slightly negative surface charge, is commonly preferred for AFM studies [26-28]. To facilitate the adsorption of negatively charged exosomes, the mica surface was rendered positively charged by treatment with APTES:DIPEA. Initially, AFM was used to assess the topography of both freshly cleaved mica and mica following APTES:DIPEA treatment. Representative height and peak force error images for freshly cleaved mica surface and APTES treated surface is shown in Figures 1A-1B and 1C-1D, respectively. No significant changes in surface topography were observed post-treatment.

**Figure 1.**
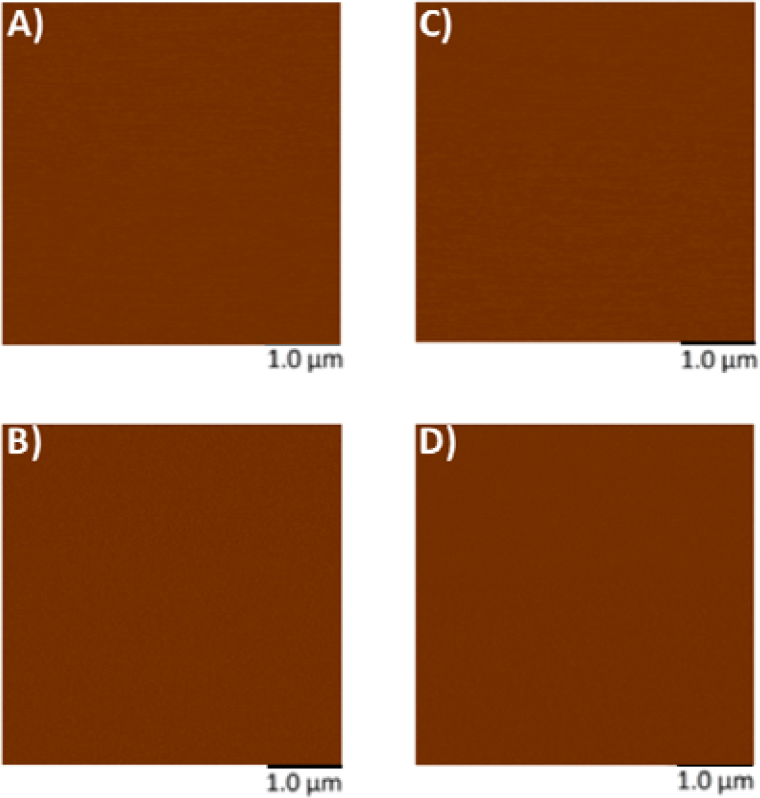
Representative surface topography of mica surface subjected to modifications. A) Height image and B)Peak force error image of **freshly cleaved mica surface**. C) Height image and DJPeak force error image of **APTES modified mica surface.**

Further, figures 2A-2B represent height image and peak force error image, respectively, of exosomes derived from Mesenchymal stem cells (MSCs). Although the peak force error image provides qualitative insights, quantitative morphological analysis was performed on the corresponding height images. The peak force error image reflects the feedback signal captured during PFQNM mode. Figures 2C-2D display height image and peak force error image, respectively, of exosomes derived from MSCs exposed to TME. o assess potential morphological differences, height images from both TME-exposed and non-exposed MSC-derived exosomes were analyzed to quantify average surface roughness and height profiles. These experiments were repeated independently on three occasions, and additional representative data are included in Supplementary Figure S1. AFM morphology imaging was conducted using probes with sharp tips, minimizing the contact area and enabling detailed resolution of surface features, as supported by previous literature [33, 34]. A reduced interaction area allows more precise detection of sudden topographical variations, yielding high-definition morphological data [25]. The topographical maps are obtained in a raster scanning format. Exosomes derived from MSCs not exposed to the TME were found to be significantly larger in size (63.28□±□4.05□nm) compared to those from TME-exposed MSCs (52.66□±□4.90□nm), as shown in Figure 2E. Conversely, MSC-derived exosomes not exposed to the TME exhibited significantly lower topographical variation (3.22□±□0.51□nm) relative to TME-exposed exosomes (5.62□±□0.32□nm), as illustrated in Figure 2F. In other words, exosomes from unexposed MSCs appeared to have smoother surfaces, whereas TME-exposed exosomes displayed increased surface heterogeneity.

**Figure 2.**
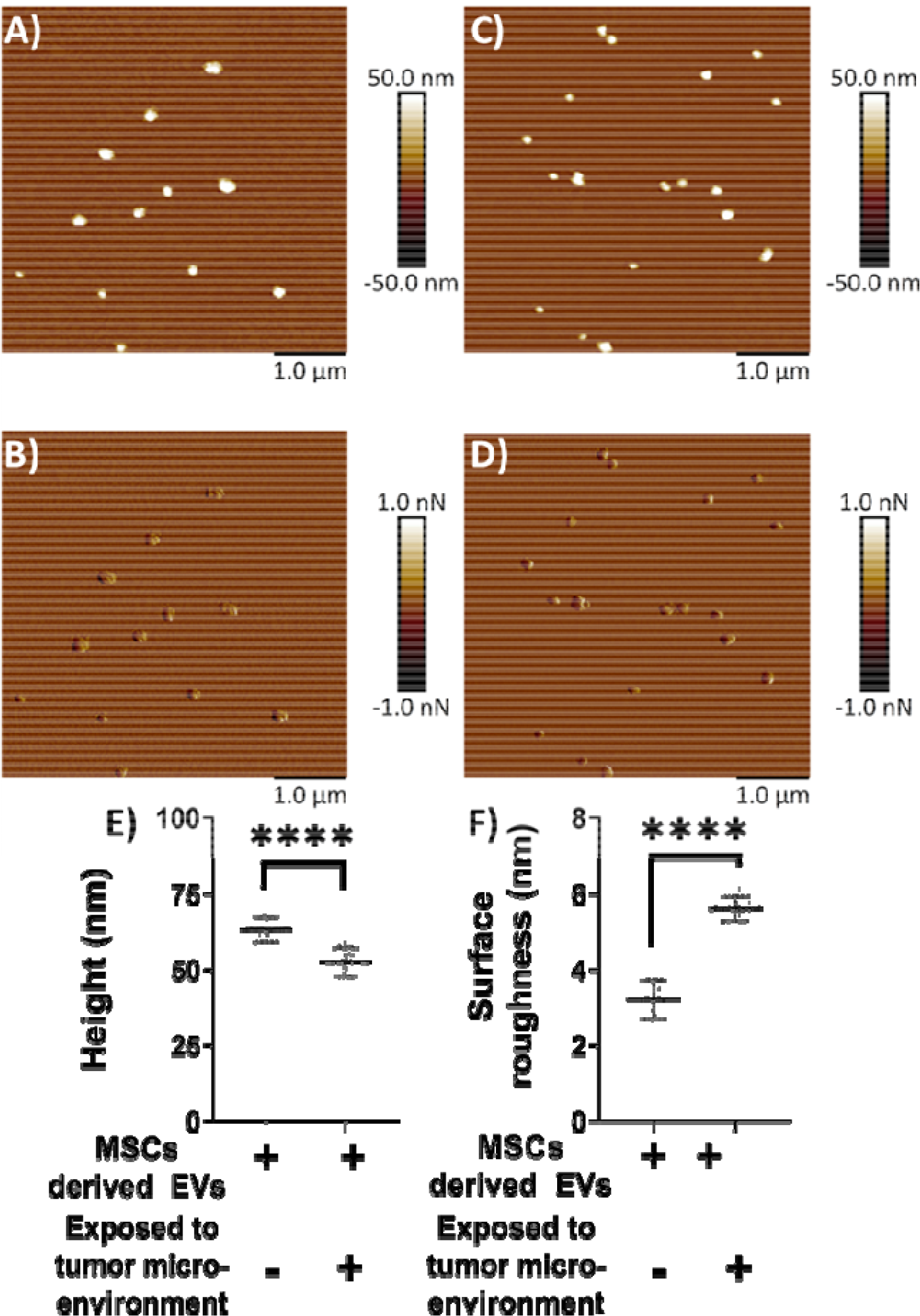
Representative surface topography of mesenchymal stem cells (MSC) derived exosomes under various conditions. A) Height image and B)Peak force error image of MSC derived exosomes non-exposed to TME. C) Height image and D)Peak force error image of MSC derived exosomes exposed to TME. Morphology quantifications. E) Height. F) Surface roughness. (Statistical significance performed by One Way ANNOVA: ****, p<0.0001)

### Nanomechanical Characterization of MSC-Derived Exosomes Under TME and Non-TME Conditions

We next focused on evaluating the nanomechanical properties of exosomes derived from MSCs under different conditions. Nanoindentation experiments were performed on exosomes obtained from both TME-exposed and non-exposed MSCs. In these experiments, each interaction between the AFM tip and the exosome surface generated a force–separation (F–S) curve. Representative F–S curves for non-exposed and TME-exposed MSC-derived exosomes are shown in Figures 3A and 3B, respectively. A rigorous optimization process was undertaken to determine the most informative ramping parameters, based on our previous studies [22-25]. The accuracy of nanomechanical measurements depends on two interrelated factors: (i) the quality and reliability of the F–S curve, and (ii) ensuring that the indentation depth remains within 20% of the exosome height to prevent excessive deformation [35]. A reliable F–S curve is characterized by a nearly flat baseline, representing negligible force during both the approach (trace) and retraction (retrace) phases when the tip is not in contact with the sample. This non-contact region is used to correct for baseline offsets.

**Figure 3.**
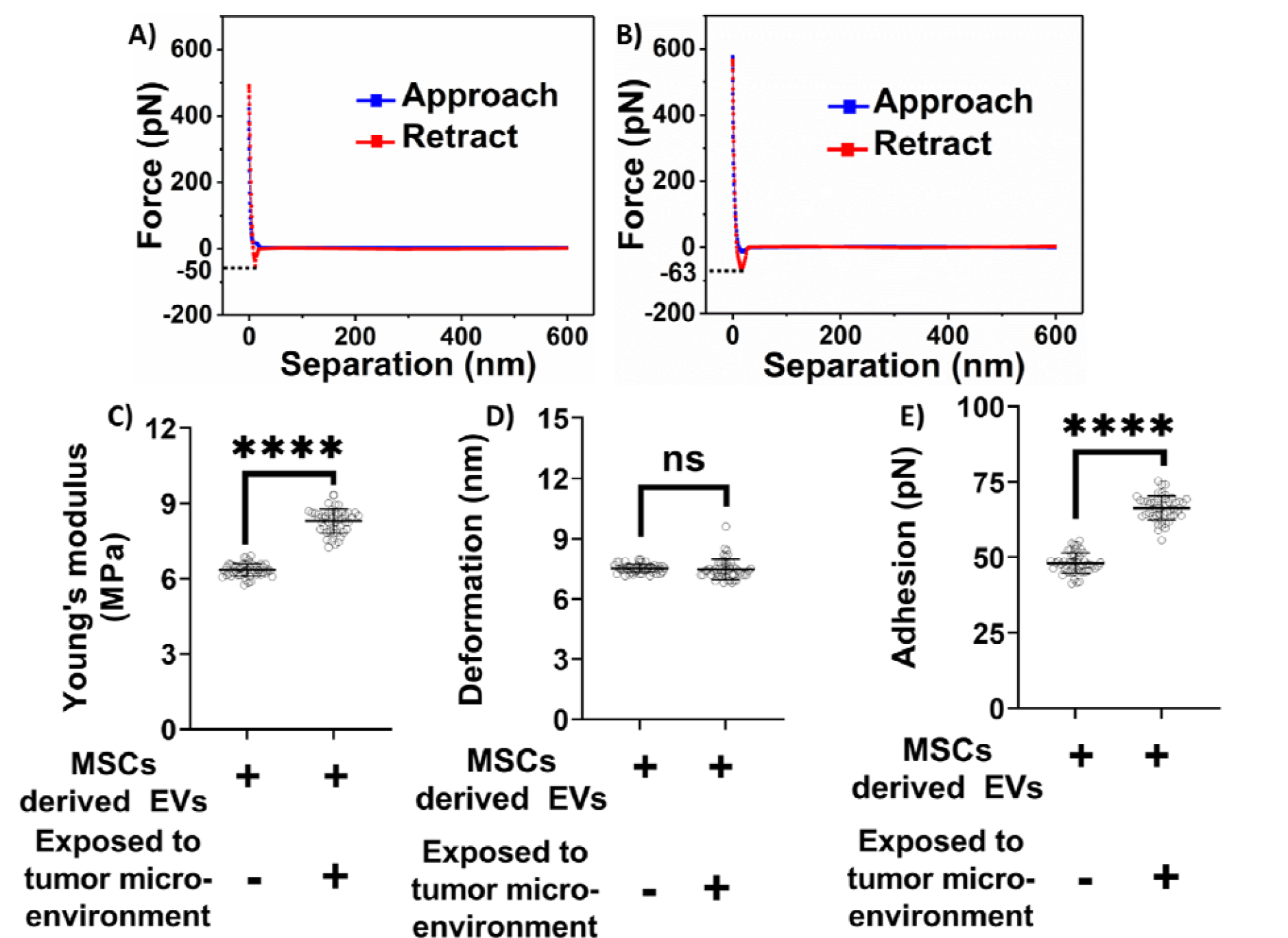
Nanomechanical characterization of mesenchymal stem cell (MSC) derived exosomes under various treatments. A representative force-separation curve for exosomes derived from MSCs and A) Not exposed to tumor microenvironment. B) Exposed to tumor microenvironment. Nanomechanical attributes C) Young’s modulus. D) Deformation. E) Adhesion. (Statistical significance performed by One Way ANNOVA: ns, not significant; ****, p<0.0001).

Based on the indentation analysis model, trace and retrace curves are used to extract Young’s modulus in various analysis models, herein Derjaguin-Muller-Toporov (DMT) model. Deformation parameter is of prime importance and is determined based on the distance between the points corresponding to slope change in trace curve and the peak force. When the tip interacts with the exosome sample during trace curve, it experiences an attractive pull towards the exosome sample and the tip begins indenting. Thereupon, the tip continues to indent till it reaches a peak user defined force, indicated by an increasing force in the trace curve. Following which, the tip starts experiencing retractive force exerted by the piezo, indicated by a decreasing force value corresponding to the retractive curve. Adhesion parameter is the result of the attractive pull experience by the tip as it retracts from the exosome surface and hence, measured from the retrace curve.

As illustrated in Figures 3A–3B, the blue line represents the approach (trace) curve, where the tip moves toward the sample, while the red line denotes the retraction (retrace) curve. Quantitative nanomechanical analysis was performed by fitting the retrace curve using the DMT model. As shown in Figure 3C, exosomes derived from MSCs not exposed to the TME were significantly softer compared to their TME-exposed counterparts. Specifically, the Young’s modulus of non-exposed exosomes was measured at 6.34□x00B1;□0.25□MPa, whereas TME-exposed exosomes exhibited a higher modulus of 8.29□±□0.47□MPa. No statistically significant difference in deformation was observed between the two groups. The deformation values were 7.51□±;□0.21□nm for non-exposed exosomes and 7.46□±□0.50□□nm for TME-exposed exosomes (Figure 3D). However, significant differences were noted in adhesion forces. Exosomes derived from MSCs not exposed to the TME exhibited an average adhesion force of 48.12□±;□3.38□pN, which was considerably lower than that of TME-exposed exosomes, which measured 66.36□±;□4.03□pN (Figure 3E). Overall, these distinct nanomechanical — especially differences in stiffness and adhesion—highlight potential biophysical markers to distinguish exosomes derived from MSCs exposed to the TME from those that are not.

## Discussion

In this study, we employed AFM to assess the morphological and nanomechanical differences between exosomes derived from MSCs exposed and not exposed to the TME. Our findings reveal that exposure to the TME significantly alters the size, surface roughness, stiffness, and adhesion characteristics of MSC-derived exosomes, suggesting that TME conditions dynamically reprogram exosomal membrane properties.

MSC-derived exosomes are increasingly recognized as key mediators within the TME, possessing both pro-tumorigenic and anti-tumorigenic properties. Their function is highly context-dependent and modulated by various molecular and cellular cues within the tumor milieu [36]. This duality renders MSC-derived exosomes both a therapeutic target and a potential tool in drug delivery and regenerative applications [37, 38].

Our morphological analyses demonstrated that TME-exposed exosomes are significantly smaller and exhibit greater topographical variation compared to non-exposed exosomes. This surface heterogeneity may reflect changes in lipid composition, protein cargo, or membrane-associated interactions, all of which can be influenced by environmental stressors within the TME[39, 40]. The observed increase in stiffness and adhesion among TME-exposed exosomes further supports this, indicating that not only the biochemical makeup but also the physical state of exosomal membranes is altered under pathological conditions.

AFM played a central role in these insights. Unlike electron microscopy, which often necessitates dehydration and staining, AFM enables the imaging of biological samples in aqueous environments, preserving native ultrastructure and avoiding preparation-induced artifacts [21]. Its nanoscale resolution and capability to probe mechanical properties—such as stiffness, elasticity, and viscoelasticity—make it an indispensable tool in membrane biology. Indeed, AFM has been instrumental in studying organelle membranes, including mitochondrial cristae, lysosomes during autophagy, and the nuclear envelope during mechanical stress or disease progression [41, 42]. The ability of AFM to measure interaction forces and monitor membrane remodeling events in real-time offers a unique advantage for understanding how exosomal membranes adapt or malfunction in disease states.

Furthermore, the differences in nanomechanical parameters—especially Young’s modulus and adhesion force—highlight their potential as physical biomarkers for differentiating exosome subtypes. These biophysical markers could be integrated into clinical workflows to identify disease-associated vesicles or assess therapeutic efficacy, especially in cancers where exosomes play a critical role in tumor progression, immune evasion, and metastasis [43, 44].

Our findings also have important implications for exosome-based drug delivery systems. The natural lipid bilayer of exosomes resembles the plasma membrane, making them highly suitable for encapsulating both hydrophobic and hydrophilic drugs [41]. The changes in membrane properties observed through AFM may influence drug loading efficiency, cargo stability, and cellular uptake—factors critical to therapeutic success. Recent studies have demonstrated that exosomes, particularly those derived from MSCs, can be engineered to express surface ligands that improve targeting to specific cell types, enhancing the efficacy of treatment in diseases like glioblastoma and Alzheimer’s [45-47]. The biophysical characterization of exosomes using AFM could thus guide the optimization of exosome production and engineering by providing real-time data on membrane integrity and surface characteristics.

However, despite the promise of exosome-based therapies, challenges remain in standardizing exosome production, improving drug-loading protocols, and ensuring reproducibility across clinical settings [42, 48]. Our AFM findings may contribute to overcoming these hurdles by offering a complementary, label-free method for assessing exosome quality and consistency.

## Conclusion

This study provides compelling evidence that the tumor microenvironment significantly influences the morphology and nanomechanical properties of MSC-derived exosomes. By leveraging the high-resolution capabilities of AFM, we identified measurable differences in exosome size, surface roughness, stiffness, and adhesion under TME conditions. These alterations not only reflect the biophysical adaptations of exosomes within a tumor context but also underscore their potential as diagnostic and therapeutic tools. Furthermore, this work highlights the broader application of AFM in studying membrane-bound structures, contributing to the evolving field of exosome engineering and targeted drug delivery.

## Acknowledgement

This research was supported by NIH-MARC Grant# 5 T34 GM100831 to Dr. H. Banerjee, DOE HBCU and NSF NOYCE Graduate training grant to Elizabeth City State University and A University of North Carolina General Administration Collaboratory research award to Dr. H. Banerjee.

---

**Fig. 1.**
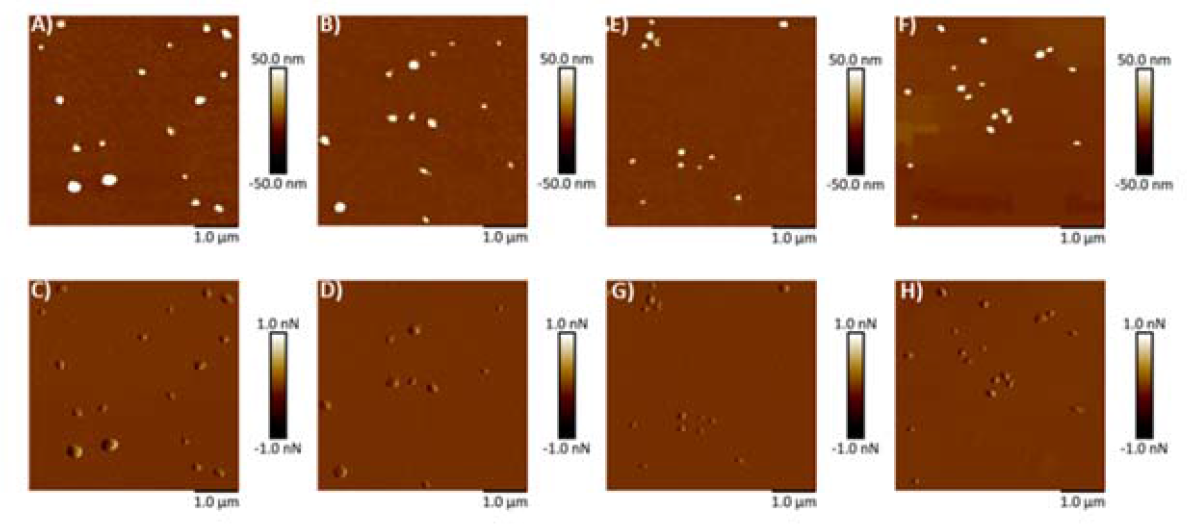
Representative surface topography images of mesenchymal stem cells derived exosomes under various conditions. A, B) Height image and C, D)Peak force error image of exosomes non-exposed to tumor microenvironment (TME). E, F) Height image and G, H) Peak force error image of exosomes exposed to TME.

## Notes

### Competing Interest Statement

The authors have declared no competing interest.

